# Transcriptional profiling of rare acantholytic disorders suggests common mechanisms of pathogenesis

**DOI:** 10.1101/2022.12.02.518412

**Authors:** Quinn R. Roth-Carter, Hope E. Burks, Ziyou Ren, Jennifer L. Koetsier, Lam C. Tsoi, Paul W. Harms, Xianying Xing, Joseph Kirma, Robert M. Harmon, Lisa M. Godsel, Abbey L. Perl, Johann E. Gudjonsson, Kathleen J. Green

## Abstract

**Background:** Darier, Hailey-Hailey, and Grover’s diseases are rare non-autoimmune acantholytic skin diseases. While these diseases have different underlying causes, they share defects in cell-cell adhesion in the epidermis and desmosome organization.

**Objective:** To better understand the underlying mechanisms leading to disease in these conditions we performed RNA-seq on lesional skin samples from Darier, Hailey-Hailey, and Grover’s disease patients.

**Methods:** RNA-seq and bioinformatics analyses were performed on banked paraffin embedded diagnostic samples from each disease. For detailed Methods, please see the Methods section in this article’s Online Repository at www.jacionline.org.

**Results:** The transcriptomic profiles of Darier, Hailey-Hailey, and Grover’s disease were found to share a remarkable overlap, which did not extend to other common inflammatory skin diseases, psoriasis and atopic dermatitis. Analysis of enriched pathways showed a shared upregulation in keratinocyte differentiation and Th17 inflammatory pathways, and a decrease in cell adhesion and actin organization pathways in Darier, Hailey-Hailey, and Grover’s disease. Direct comparison to atopic dermatitis and psoriasis showed that the downregulation in actin organization pathways was a unique feature in Darier, Hailey-Hailey, and Grover’s disease.

Further, upstream regulator analysis suggests that a decrease in SRF/MRTF activity may be responsible for the downregulation of actin organization pathways. Staining for MRTFA in lesional skin samples showed a decrease in nuclear MRTFA in patient skin compared to normal skin.

**Conclusion:** These findings highlight the significant level of similarity in the transcriptome of Darier, Hailey-Hailey, and Grover’s disease, and identify decreases in actin organization pathways as a unique signature present in these conditions.

**Key Messages:**

- Darier Disease, Hailey-Hailey Disease, and Grover’s Disease share similar transcriptional profiles suggesting common mechanisms of pathogenesis.
- SRF/MRTFA activity is reduced in Darier Disease, Hailey-Hailey Disease and Grover’s disease, implicating actin organization in acantholysis.

## Introduction

Acantholysis, or loss of adhesion between keratinocytes, is a common feature shared by the non-autoimmune skin diseases, Darier disease (DD), Hailey-Hailey disease (HHD), and Grover’s disease (GD). While these diseases have varying clinical presentations and etiologies, they share acantholysis, and loss of desmosome function as major features. DD and HHD are caused by mutations in the calcium channels ATP2A2 or ATP2C1 respectively, suggesting that calcium dysregulation is a shared feature between DD and HHD, while the etiology for GD is unknown (1, 2). The biological mechanisms leading to disease in these conditions is largely unknown, limiting the clinical strategies for treatment to symptom reduction.

To better characterize the shared and divergent molecular and cellular processes driving DD, HHD and GD we performed transcriptome profiling on lesional skin samples. The results revealed that the transcriptional profiles of DD, HHD and GD are more similar to each other than to the common inflammatory skin conditions atopic dermatitis (AD) and psoriasis (PSO). Pathway analysis revealed unique signatures in DD, HHD and GD, in particular a downregulation in actin organization pathways, which may highlight novel underlying mechanisms leading to disease in these patients.

## Results and Discussion

### Transcriptome profiling of DD, HHD and GD patient samples reveals high level of similarity

Principle component analysis (PCA) of RNA-seq results from DD, HHD and GD showed a clustering of disease samples away from controls, however; the disease samples formed a larger mixed cluster (Figure 1A). Comparison of upregulated and downregulated genes showed substantial overlap in the genes across conditions (Figure 1B, C). Spearman correlations between disease conditions revealed a significant overlap among all three conditions (Figure 1D-F). Based on these observations we tested if pathways known to be upregulated in DD, such as pathways associated with ER stress, were also present in the other conditions (3, 4). We performed upstream regulator analysis using Ingenuity Pathway Analysis (IPA) to determine if transcription factors that regulate ER stress response genes were predicted to be activated and found a significant increase in predicted ATF4 activity compared with control samples, and an upregulation in expression of many ATF4 target genes in DD, HHD and GD (Figure 1G). These observations suggest that while these diseases do have different etiology and presentation, the underlying changes in the skin transcriptome are similar.

**Figure 1.**
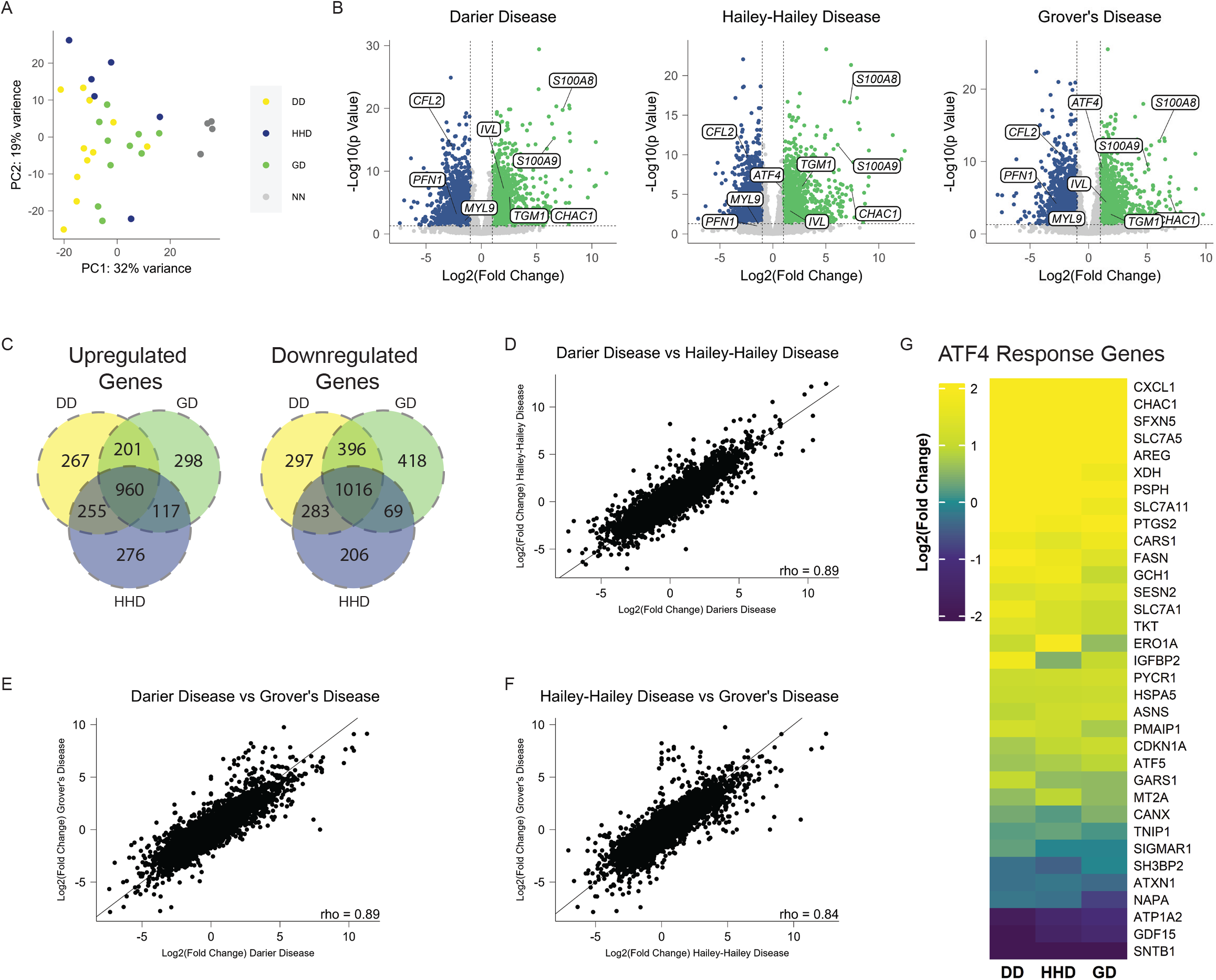
Whole transcriptome profiling of Darier, Hailey-Hailey and Grover’s disease samples reveals high level of similarity between conditions. A) Principal component analysis of samples from DD, HHD, GD, and NN. B) Volcano plot showing significantly upregulated (green) and downregulated (blue) genes in DD, HHD and GD compared to NN skin. C) Venn diagram showing overlap in significantly changed genes in DD, HHD, and GD. D-F) Correlation analysis of gene expression values from DD, HHD and GD. G) Heatmap showing gene expression of ATF4 response genes.

### DD, HHD and GD share changes in keratinocyte differentiation and cell-cell adhesion pathways

Gene Set Enrichment Analysis (GSEA) on all conditions using Gene Ontology (GO) Biological Process (BP) pathways revealed increases in pathways associated with keratinocyte differentiation and decreases in pathways associated with cell-cell adhesion, actin organization, and RHO signaling (Figure 2A). To further explore the changes in keratinocyte differentiation we used single-cell RNA-seq data collected from normal skin to create gene set signatures using genes expressed specifically in basal, differentiated, and keratinized keratinocytes (5).

**Figure 2.**
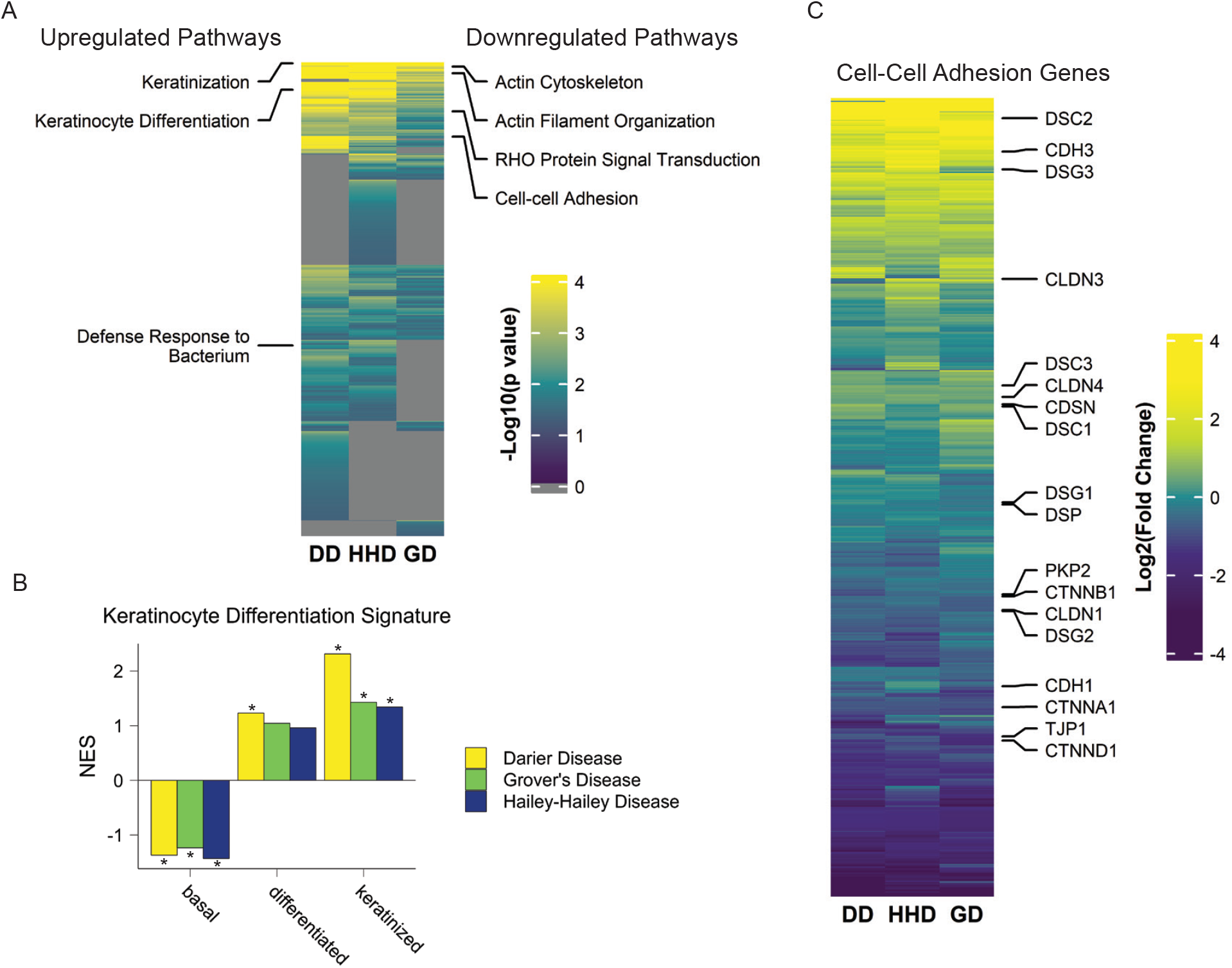
Pathway analysis reveals shared changes in keratinocyte differentiation and cell-cell adhesion in DD, HHD and GD. A) Heatmap showing GO BP pathways ranked by -Log10(p value) for DD, HHD and GD. Selected upregulated pathways are noted on the left of the heatmap, while downregulated pathways are noted on the right. B) DD, HHD and GD gene expression changes were compared to keratinocyte differentiation signatures using GSEA. *FDR < 0.05. C) Heatmap showing log_2_(FoldChange) values compared with controls for genes annotated to the GO BP cell-cell adhesion pathway.

Comparison of these gene sets to DD, HHD and GD using GSEA revealed a decrease in basal associated genes, and a significant increase in keratinized associated genes in all conditions, suggesting a defect in normal keratinocyte differentiation (Figure 2B). We also observed a decrease in pathways associated with cell-cell adhesion, fitting with observations of acantholysis in the skin in these patients (Figure 2C).

### DD, HHD and GD share greater similarities to each other than to psoriasis and atopic dermatitis

We next compared DD, HHD and GD to other more common skin diseases, PSO and AD, to determine shared and unique features across these diseases. To perform direct comparisons between each condition the DD, HHD and GD samples were pooled with a publicly available PSO and AD dataset, and batch corrected (6). Spearman correlation followed by hierarchical clustering showed that PSO, AD, and control samples tend to cluster together, while DD, HHD, and GD formed two intermixed clusters (Figure 3A). Similarly, dimensional reduction using UMAP showed AD, PSO, and controls separating into distinct clusters, while DD, HHD, and GD cluster into a mixed group (Figure 3B)(7), suggesting that these diseases share greater similarity to each other than to AD or PSO. Comparing overlap in significantly upregulated and downregulated genes between PSO, AD, and combined non-autoimmune acantholytic disease samples showed significant overlap in upregulated genes across all disease, but non-significant overlap in downregulated genes between the non-autoimmune acantholytic diseases and PSO and AD, suggesting that the difference in transcriptional signatures is largely driven by downregulated genes in DD, HHD and GD (Figure 3C). GSEA using GO BP pathways revealed shared overlap in upregulated pathways involved in keratinocyte differentiation and inflammatory responses between DD, HHD, GD, PSO and AD (Figure 3D). When analyzing downregulated pathways, we observed numerous pathways that were only downregulated in DD, HHD, and GD but not AD and PSO, such as actin cytoskeleton organization and focal adhesion assembly (Figure 3E). While these non-autoimmune acantholytic skin diseases are typically associated with desmosome dysfunction, these observations suggest that there is actin cytoskeleton dysregulation as well (8-11).

**Figure 3.**
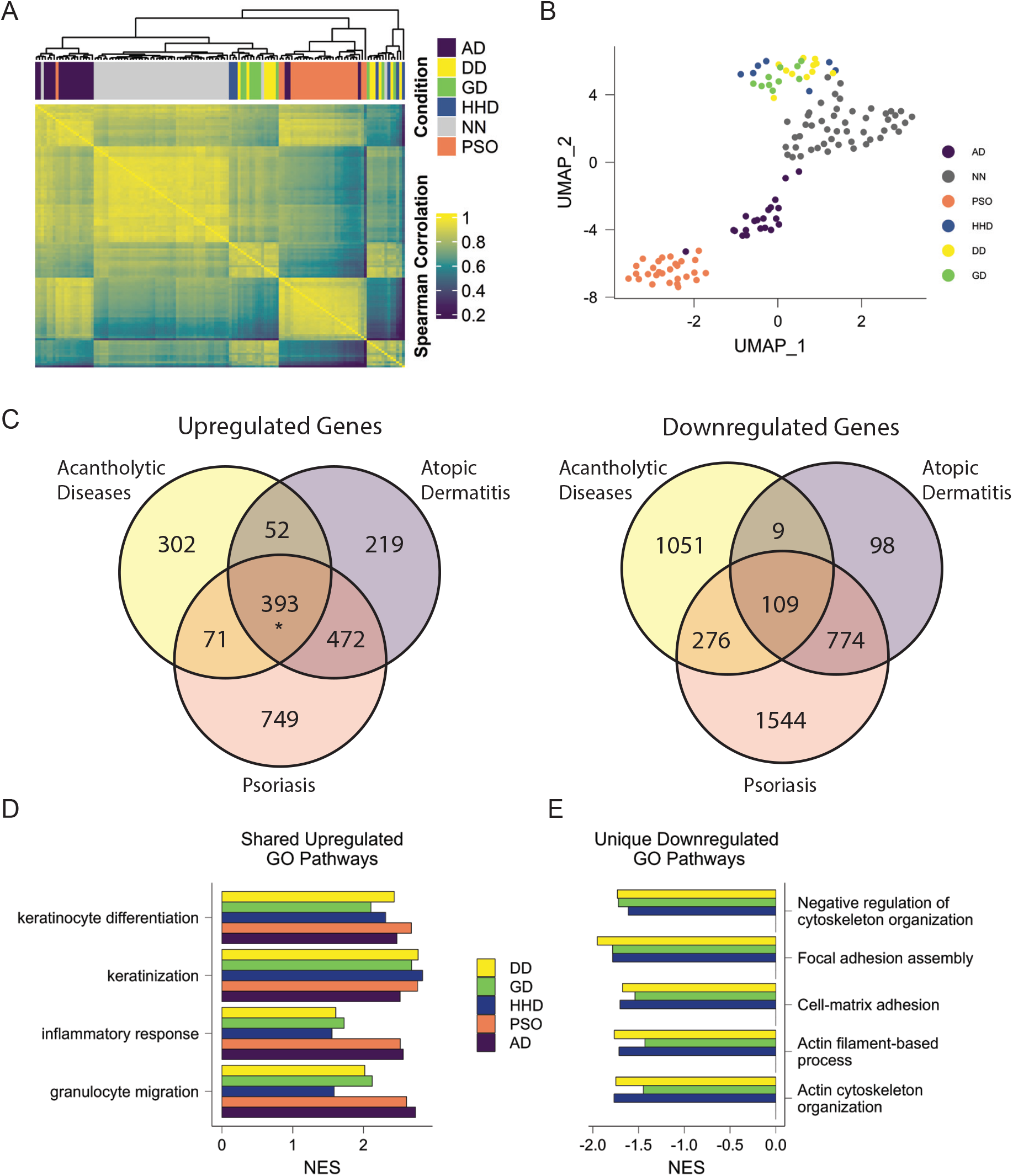
DD, HHD and GD are more similar to each other than to PSO and AD. A) Heatmap showing Spearman correlation values between the batch-adjusted counts from all individual samples grouped using hierarchical clustering. B) Dimensional reduction plot using UMAP on all individual samples. C) Venn diagrams showing overlap in genes upregulated or downregulated in PSO, AD, and combined acantholytic diseases (p=0.4 for downregulated genes and p<0.00001 for upregulated genes by permutation test). D) Significantly upregulated GO BP pathways present in all conditions. E) GO BP pathways significantly downregulated in DD, HHD and GD, but unchanged in PSO and AD.

### DD, HHD and GD share a weak Th17 inflammatory signature that is similar to psoriasis

Previous work with the AD and PSO data sets has already shown that IL-17 and IFNG response pathways are upregulated in both, while IL-13 response pathways are upregulated specifically in AD (6). However, the inflammatory signatures associated with DD, HHD, and GD are unknown; therefore, we used IPA to examine predicted cytokine upstream regulators. We observed an increase in IFNG, IL-17A, IL-36G and IL-36A responses in all conditions (Figure 4A). However, the extent of enrichment was lower in DD, HHD and GD compared to AD and PSO. IL-13 responses were only enriched in AD samples (Figure 4A). Analyzing gene expression of defining cytokines in these response pathways showed modest statistically significant increases in expression of IL17A in DD and HHD, while other cytokines were not upregulated in these conditions (Figure 4B, C). These data suggest that DD, HHD and GD share a Th17 inflammatory signature, though one that is not as prominent as what is observed in PSO. These observations are in agreement with descriptions of the immune infiltrate in DD, HHD and GD, with a low level of inflammatory cell recruitment in lesional skin, and together suggest that the inflammatory signature is not a critical driver of disease (12).

**Figure 4.**
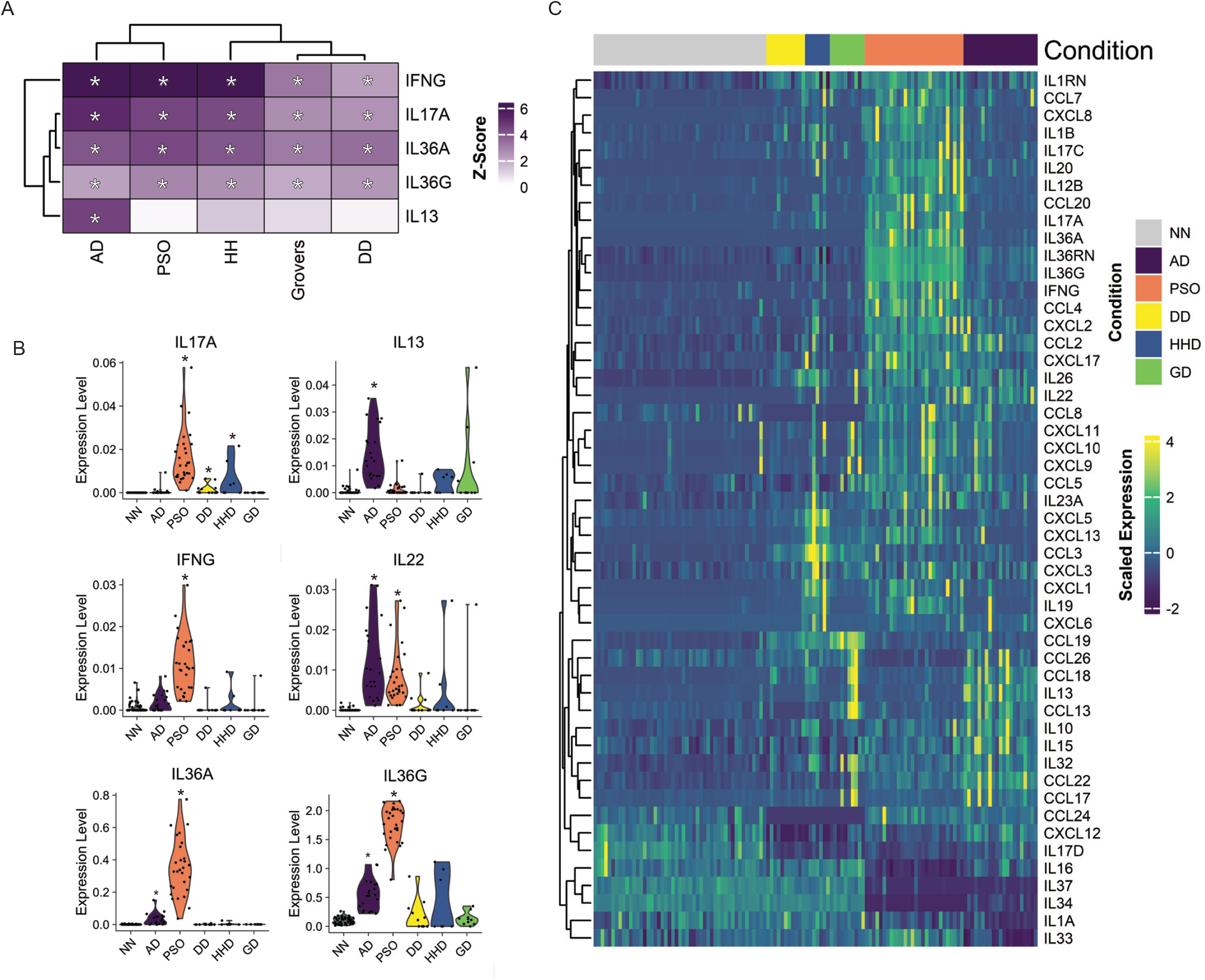
Th17 inflammatory signatures are common across DD, HHD, GD, PSO and AD, while only AD has Th2 inflammatory signatures. A) Heatmap showing z-scores of predicted upstream regulators identified using Ingenuity Pathway Analysis (IPA) with a focus on cytokines. B) Violin plots demonstrating expression levels of 6 cytokines from all conditions including controls (NN). C) Heatmap depicting scaled expression of cytokines and chemokines across conditions.

### SRF/MRTF activity is predicted to be downregulated in the skin in DD, HHD, and GD patients

We next sought to examine unique features present in the non-autoimmune acantholytic skin disease compared to PSO and AD. The downregulation of actin organization pathways was identified as a distinct feature in DD, HHD and GD compared to AD and PSO (Figure 3E). To explore factors responsible for the downregulation of actin organization pathways in DD, HHD and GD we focused on two major known transcription factors that regulate actin organization: serum response factor (SRF) and yes-associated protein/transcriptional coactivator with PDZ-binding motif (YAP/TAZ). Upstream regulator analysis revealed a predicted decrease in SRF activity, with no change in YAP/TAZ in DD, HHD, and GD (Figure 5A), indicating that a reduction in SRF activity may be responsible for the observed downregulation of actin organization pathways in these conditions. GSEA of predicted transcription factor target genes using the transcription factor targets database from MSigDB revealed a decrease in genes containing SRF binding motifs, with a trend towards an increase in TAZ target genes (Figure 5B) (13-15). This observation is in line with the upstream regulator findings, again suggesting that SRF activity is reduced in the skin of patients with DD, HHD and GD. Additionally, we assessed changes in SRF cofactors. The primary mechanisms by which SRF can influence gene expression are through interaction with MRTFs or interactions with the ternary complex factors (ELK-1, ELK-3, and ELK-4). Upstream regulator analysis predicts decreased activity of MRTFA and MRTFB, while activity of ELK-1 ELK-3 and ELK-4 remains unchanged in DD, HHD and GD (Figure 5C). Finally, to validate these observations we stained patient skin samples for MRTFA and YAP1. We found that the nuclear/cytoplasmic staining intensity for MRTFA was significantly reduced in patient samples, while YAP staining, though highly variable, showed a trend towards greater levels in patient samples. (Figure 5D, E). Given the importance of SRF/MRTFA signaling in epidermal differentiation and barrier formation, its dysregulation in these disorders is likely a contributor to pathogenesis of these diseases (16).

**Figure 5.**
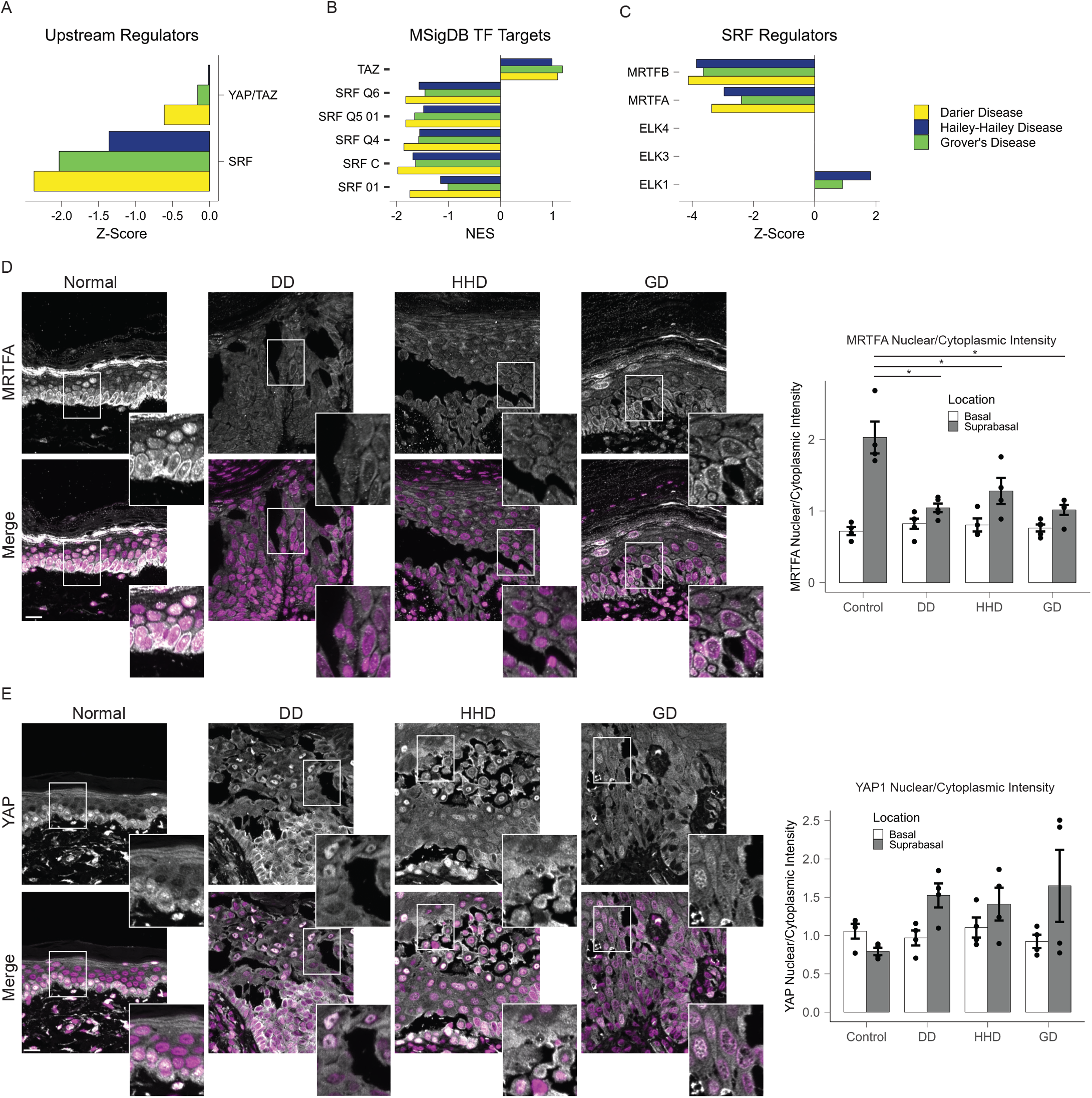
SRF is predicted to be downregulated in acantholytic skin diseases. A) Predicted activity of actin organization regulating transcription factors identified using upstream regulator analysis. B) Enrichment (NES) of SRF target sequences and TAZ target genes from MSigDB predicted TF Targets database. C) IPA upstream regulator analysis showing predicted activity of SRF cofactors. Signatures were not detected for ELK3 and 4. D) Immunostaining for MRTFA in DD, HHD, GD and NN skin. Quantification of MRTFA nuclear and cytoplasmic pixel intensities measured in basal and suprabasal cells. Scale bar = 20µm. E) Immunostaining for YAP1 in DD, HHD, GD and NN skin. Quantification of YAP1 nuclear and cytoplasmic pixel intensities measured in basal and suprabasal cells. Scale bar = 20µm.

The acantholysis seen in these conditions is commonly attributed to desmosome dysfunction. While desmosomes are classically defined as organizers of the intermediate filament cytoskeleton, previous work from our group has demonstrated that desmosomes can regulate actin remodeling (17-19). Collectively, these observations raise the possibility that acantholysis in patients with DD, HHD and GD, is driven by interference with desmosomes and associated actin organization. Furthermore, this study is the first to mechanistically link GD with characterized calcium pump mutation-driven acantholytic disorders. While the question of calcium signaling dysregulation in GD remains, the downstream transcriptional profiles share remarkable overlap in all conditions and present novel regulatory pathways at play in poorly understood non-autoimmune acantholytic diseases.

## Supporting information

Online Repository

## Abbreviations

AD: Atopic Dermatitis
DD: Darier Disease
ELK1: ETS transcription factor ELK1
ELK3: ETS transcription factor ELK3
ELK4: ETS transcription factor ELK4
HHD: Hailey-Hailey Disease
IPA: Ingenuity Pathway Analysis
GD: Grover’s Disease
GSEA: Gene Set Enrichment Analysis
MRTFA: Myocardin-related transcription factor A
MRTFB: Myocardin-related transcription factor B
PCA: Principle Component Analysis
PSO: Psoriasis
SRF: Serum response factor
TAZ: transcriptional coactivator with PDZ-binding motif
YAP: Yes-associated protein

## Acknowledgments

We thank the members of the Green Lab for their valuable feedback and discussion. We also thank the Lee family for their generous support.

## References

1. Sakuntabhai A, Ruiz-Perez V, Carter S, Jacobsen N, Burge S, Monk S, et al. Mutations in ATP2A2, encoding a Ca2+ pump, cause Darier disease. Nature Genetics. 1999;21(3):271–7.

2. Sudbrak R, Brown J, Dobson-Stone C, Carter S, Ramser J, White J, et al. Hailey–Hailey disease is caused by mutations in ATP2C1 encoding a novel Ca2+ pump. Human Molecular Genetics. 2000;9(7):1131–40.

3. Onozuka T, Sawamura D, Goto M, Yokota K, Shimizu H. Possible role of endoplasmic reticulum stress in the pathogenesis of Darier’s disease. Journal of Dermatological Science. 2006;41(3):217–20.

4. Savignac M, Simon M, Edir A, Guibbal L, Hovnanian A. SERCA2 dysfunction in Darier disease causes endoplasmic reticulum stress and impaired cell-to-cell adhesion strength: rescue by Miglustat. J Invest Dermatol. 2014;134(7):1961–70.

5. Godsel LM, Roth-Carter QR, Koetsier JL, Tsoi LC, Huffine AL, Broussard JA, et al. Translational implications of Th17-skewed inflammation due to genetic deficiency of a cadherin stress sensor. J Clin Invest. 2022;132(3).

6. Tsoi LC, Rodriguez E, Degenhardt F, Baurecht H, Wehkamp U, Volks N, et al. Atopic Dermatitis Is an IL-13-Dominant Disease with Greater Molecular Heterogeneity Compared to Psoriasis. J Invest Dermatol. 2019;139(7):1480–9.

7. McInnes L, Healy J, Melville J. Umap: Uniform manifold approximation and projection for dimension reduction. arXiv preprint 180203426. 2018.

8. Hashimoto K, Fujiwara K, Tada J, Harada M, Setoyama M, Eto H. Desmosomal dissolution in Grover’s disease, Hailey-Hailey’s disease and Darier’s disease. J Cutan Pathol. 1995;22(6):488–501.

9. Hashimoto K, Fujiwara K, Harada M, Setoyama M, Eto H. Junctional proteins of keratinocytes in Grover’s disease, Hailey-Hailey’s disease and Darier’s disease. J Dermatol. 1995;22(3):159–70.

10. Hobbs RP, Amargo EV, Somasundaram A, Simpson CL, Prakriya M, Denning MF, et al. The calcium ATPase SERCA2 regulates desmoplakin dynamics and intercellular adhesive strength through modulation of PKCα signaling. Faseb j. 2011;25(3):990–1001.

11. Hakuno M, Shimizu H, Akiyama M, Amagai M, Wahl JK, Wheelock MJ, et al. Dissociation of intra- and extracellular domains of desmosomal cadherins and E-cadherin in Hailey-Hailey disease and Darier’s disease. Br J Dermatol. 2000;142(4):702–11.

12. See SHC, Peternel S, Adams D, North JP. Distinguishing histopathologic features of acantholytic dermatoses and the pattern of acantholytic hypergranulosis. Journal of Cutaneous Pathology. 2019;46(1):6–15.

13. Subramanian A, Tamayo P, Mootha VK, Mukherjee S, Ebert BL, Gillette MA, et al. Gene set enrichment analysis: A knowledge-based approach for interpreting genome-wide expression profiles. Proceedings of the National Academy of Sciences. 2005;102(43):15545–50.

14. Kolmykov S, Yevshin I, Kulyashov M, Sharipov R, Kondrakhin Y, Makeev VJ, et al. GTRD: an integrated view of transcription regulation. Nucleic Acids Res. 2021;49(D1):D104–d11.

15. Xie X, Lu J, Kulbokas EJ, Golub TR, Mootha V, Lindblad-Toh K, et al. Systematic discovery of regulatory motifs in human promoters and 3′ UTRs by comparison of several mammals. Nature. 2005;434(7031):338–45.

16. Koegel H, von Tobel L, Schäfer M, Alberti S, Kremmer E, Mauch C, et al. Loss of serum response factor in keratinocytes results in hyperproliferative skin disease in mice. J Clin Invest. 2009;119(4):899–910.

17. Broussard JA, Yang R, Huang C, Nathamgari SSP, Beese AM, Godsel LM, et al. The desmoplakin-intermediate filament linkage regulates cell mechanics. Mol Biol Cell. 2017;28(23):3156–64.

18. Godsel LM, Dubash AD, Bass-Zubek AE, Amargo EV, Klessner JL, Hobbs RP, et al. Plakophilin 2 couples actomyosin remodeling to desmosomal plaque assembly via RhoA. Mol Biol Cell. 2010;21(16):2844–59.

19. Nekrasova O, Harmon RM, Broussard JA, Koetsier JL, Godsel LM, Fitz GN, et al. Desmosomal cadherin association with Tctex-1 and cortactin-Arp2/3 drives perijunctional actin polymerization to promote keratinocyte delamination. Nature Communications. 2018;9(1):1053.

