## Supplementary material for "Transcriptional profiling of rare acantholytic disorders suggests common mechanisms of pathogenesis": Online Repository

### METHODS

#### Study Approval

Darier disease, Hailey-Hailey disease, Grover's disease patients and healthy adults provided informed consent for skin biopsies. Tissues were anonymized for analysis and collected under IRB# HUM00087890 at University of Michigan.

#### Additional Gene Expression Datasets

Atopic Dermatitis and Psoriasis RNAseq dataset was downloaded from Gene Expression Omnibus (GEO), accession number GSE121212. This dataset is originally described in (1). Single-cell RNA-seq dataset used to generate keratinocyte differentiation genes sets is available in GEO, accession number GSE179162.

#### RNA-Seq expression profiling of Darier, Hailey-Hailey and Grover's disease samples.

RNA was isolated from 10 um sections of formalin-fixed paraffin embedded blocks from 11 Darier disease, 7 Hailey-Hailey disease, and 10 Grover's disease samples. RNA was isolated using the E.N.Z.A. FFPE RNA Kit (Omega Bio-tek). Samples were prepared using the Lexogen 3' QuantSeq mRNA-Seq Library Prep Kit FWD and sequences on the Illumina NovaSeq 6000 System. Quality control and adaptor trimming were performed on sequence reads form the RNA-Seq data. STAR alignment was used to align the reads to the reference genome (GRCh37), and HTSeq was used for gene quantification. To eliminate potential differences caused by sex specific genes, Y chromosome genes and genes known to be differentially expressed between males and females in skin were removed from further analysis (2). To generate differential expression for Darier, Hailey-Hailey, and Grover's disease the DESeq2 Bioconductor R Package V1.34.0 was used.

#### Correlation Analysis between Darier disease, Hailey-Hailey disease, Grover's disease, Psoriasis and Atopic Dermatitis.

Analysis was performed in R V4.1.1 (R Foundation for Statistical Computing). Batch correction was performed using ComBat-seq function under the SVA Bioconductor R package V3.36.0.0 following published method (3). PCA was computed based on batch-adjusted raw counts using R package pcaMethods V1.86.0. For comparisons between Darier Disease, Hailey-Hailey Disease, Grover's disease, psoriasis, and atopic dermatitis the EdgeR Bioconductor R Package V3.36.0 was used to generate differential expression data from batch-adjusted raw counts and calculate statistically significantly changed genes by ANOVA. Genes that had an FDR-adjusted p value < 0.05 and $|Log_2(\text{Fold Change})| > 1$ in any one condition were used for all further comparison analysis. Spearman correlations were calculated using the batch-adjusted raw counts and the *cor* function in R build-in stats package. UMAP plots were created using the Seurat package in R V4.1.1.

#### Functional Enrichment Analysis

Functional enrichment analysis was performed using the clusterProfiler package in R V4.2.2 and using Ingenuity Pathway Analysis (IPA) software. Pathway analysis was performed using Gene Ontology Biologic Process and the Transcription Factor Targets pathways, and were analyzed using Gene Set Enrichment Analysis (GSEA)(4).

### Immunofluorescence and image acquisition

DD, HHD, GD and normal patient skin samples were fixed in 10% formalin, embedded in paraffin blocks and cut into 4μm thick sections. For immunostaining paraffin sections were baked at 60°C overnight and de-paraffinized with xylene. Samples were then rehydrated through a series of ethanol and PBS washes, and permeabilized in 0.5% Triton X-100 in PBS. Antigen Retrieval was performed by incubating the slides in 0.01 M citrate buffer at 95°C for 15 minutes. Samples were blocked in blocking buffer (1% BSA, 2% normal goat serum in PBS) for 1 hour at 37°C. Samples were then incubated in primary antibody overnight at 4°C, followed by washes with PBS and incubation in secondary antibody for 1 hour at 37°C. Images were acquired using an AxioVision Z1 system (Carl Zeiss) with an Apotome slide module, and AxioCam MRm digital camera and a 20x (0.8 NA Plan-Apochromat) objective. Image analysis was performed using ImageJ software.

### Antibodies

Antibodies used in this study include: rabbit anti-YAP1 (14074, Cell Signaling), rabbit anti-MRTFA (PA599446, ThermoFisher Scientific), AlexaFluor 568-conjugated goat anti-rabbit secondary antibodies (ThermFisher Scientific).

### Statistics Analysis

Permutation analysis for Venn Diagrams was performed to test the significance of the overlap. 10,000 permutations were conducted from non-replacement random sampling of 6,000 tokens. An empirical p value is calculated by comparing the actual and the permutation results.

For image analysis statistical analysis was performed using one-way ANOVA with Dunnett correction for multiple comparisons, with all disease samples compared only to control samples. $P < 0.05$ was considered statistically significant, and data represent mean ± SEM.

### Data Availability

The resulting RNA-seq data sets generated for this study will be available in GEO at the time of publication, and all code used to analyze the data will be available.

### References


1. Tsoi LC, Rodriguez E, Degenhardt F, Baurecht H, Wehkamp U, Volks N, et al. Atopic Dermatitis Is an IL-13-Dominant Disease with Greater Molecular Heterogeneity Compared to Psoriasis. J Invest Dermatol. 2019;139(7):1480-9.
2. Gershoni M, Pietrokovski S. The landscape of sex-differential transcriptome and its consequent selection in human adults. BMC Biology. 2017;15(1):7.
3. Zhang Y, Parmigiani G, Johnson WE. ComBat-seq: batch effect adjustment for RNA-seq count data. NAR Genom Bioinform. 2020;2(3):lqaa078.
4. Liberzon A, Birger C, Thorvaldsdóttir H, Ghandi M, Mesirov JP, Tamayo P. The Molecular Signatures Database (MSigDB) hallmark gene set collection. Cell Syst. 2015;1(6):417-25.
